# A new method of teaching medical statistics at medical universities in Guangdong

**DOI:** 10.1101/815142

**Authors:** Qing-shan Chen, Xin-yue Ma, Xin Cheng, Xuesong Yang

## Abstract

Although statistics play a significant role in the medical profession, studying medical statistics is challenging because this topic is more difficult to comprehend than other subjects included in medical curricula. Therefore, improving the teaching of medical statistics to meet the requirements of modern medical students/physicians is an essential task. In this study, based on responses about studying medical statistics completed by medical students, we developed a novel approach to teaching medical statistics named “purpose, database, types of variable, and relationship between variables (PDTR)”, which emphasizes how to simply master statistical applications and reduces class hours for students. Also, pilot course was implemented. We discovered that the participants using PDTR performed better on examinations of medical statistics than did graduates of institutions using traditional teaching methods (86.39±3.72 vs 73.72±6.58, P < 0.001). In addition, positive feedback was received by participants (>80%). Altogether, as a completely novel pedagogical method of teaching medical statistics, PDTR overcomes the negative attitudes of students towards medical statistics, enhances enthusiasm for learning statistics, and remarkably simplifies the process of studying statistical applications. These advantages are undoubtedly conductive to improving the effective use of medical statistics in physicians’ professional work.

## Introduction

The current Chinese medical education system training physicians in Western medicine was established in Chinese medical schools based on the model of Western medicine(1), which has been developed over centuries in Western countries. Medical students, especially those in clinical medicine, must have strong medical knowledge including pre-clinical and clinical sciences such as anatomy, physiology, biochemistry, pharmacology, medicine, surgery, etc., which is certainly a considerable study pressure for them. Undoubtedly, understanding medical statistics is absolutely indispensable, even for physicians who do not engage in medical research as a part of their occupation. Medical doctors are certainly expected to be able to carry out statistical analyses in their routine work. Physicians must always consider averages and standard deviations while prescribing drugs or implementing medical practices in the clinic, integrating or scientifically evaluating biological evidence/deviations, analysing the symptoms of complicated diseases, or even discovering novel drugs and therapies; thus, medical statistics are required for many tasks. In addition, due to the rapid development of information technology and evidence-based medicine, medical statistical training for modern doctors should be updated(2) because statistics are the core basis of all quantitative research(3). Medical statistics education at Chinese medical universities/schools has been developed for several decades based on the curriculum guidelines used in Western countries(4). Unfortunately, compared to other medical subjects, the study of medical statistics is associated with great frustration and difficulty among many Chinese medical students(5); in addition, these attitudes are stronger than those identified in medical students attending Western universities(6, 7). Hence, improving the methods used to teach medical statistics in most Chinese universities/schools is becoming urgent.

All higher education curricula intend to cultivate students’ and graduates’ competences in solving problems rather than considering only the teaching outcome. Thus, the following question emerged: Are the students’ perspectives ignored in designing curricula for current medical statistics courses as in other medical educational curriculum design? (8) Mules et al. reported that medical statisticians dominate the design of statistics textbooks used to teach UK medical undergraduates without full consideration of the physicians who will actually use the statistical knowledge after graduation (2). Thus, there is a great need to explore the views of medical students and physicians regarding statistical training and curriculum, especially based on their actual requirement for statistics in daily medical practice.

Although rapid and robust achievements have been realized in the field of medical education and studies in China, the current status of medical education, including medical statistics, is still unsatisfactory and weak; in particular, progress in pedagogical theories is slow compared to that at the top medical schools worldwide (9). Moreover, the rapid development of life sciences over the last few decades has dramatically enriched medical knowledge, leading to the reform of medical education worldwide. However, to date, Chinese medical statistics programmes have generally remained unchanged partially because reform requires substantial complex work.

In this study, first, we investigated medical students’ attitudes towards the current state of medical statistics teaching at three medical universities/schools in Guangdong Province; then, an applied medical statistics approach was developed based on students’ actual requirements, and pilot courses were applied over the past few years. Subsequently, a survey evaluating this novel teaching approach was also performed.

## Material and methods

### Current status of Chinese medical statistics

As a compulsory course, medical statistics is taught to medical students during their third or fourth year in all Chinese medical universities/schools according to the syllabus requirements of the Education Ministry. Here, the syllabus of medical statistics in Jinan University School of Medicine was presented in Table 1 and Table 2. Medical Statistics course consist of two aspects: classroom teaching and practice. Classroom teaching focuses on basic statistical concepts, such as how to select statistical methods based on obtained data, how to perform SPSS statistics software, how to interpret and report the output of SPSS. The each course takes 2 class hours and total contact hours are 28 class hours. Practice teaching: the aim is to guide students on analyzing statistical data by SPSS on computers. There are eight lessons each semester, and total contact hours are 16 class hours (two hours each lesson). Therefore, there are totally 44 class hours in each semester (Note: 1 class hour = 45 minutes).

**Table 1.** Key points and class hour allocation in classroom teaching (traditional)

| Chapter | Teaching<br>hour |
| --- | --- |
| 1. Introduction | 2 |
| 2. Statistical Description of quantitative data | 4 |
| 3. Sampling Error and Estimation of Population Mean | 2 |
| 4. Hypothesis | 2 |
| 5. *t* test | 2 |
| 6. Analysis of Variance | 2 |
| 7. Relative Measures | 2 |
| 8. $\chi^2$ Test for Categorical Variable | 2 |
| 9. Nonparametric Test Based on Ranks | 2 |
| 10. Linear Regression and Correlation | 2 |
| 11. Statistical Table and Graph | 2 |
| 12. Statistical Design in Medical Research | 4 |

**Table 2.** Key points and class hour allocation in practice teaching (traditional)

| Experiments | Teaching hour |
| --- | --- |
| 1.To Start SPSS and Input Data | 2 |
| 2.Sorting and Transforming Data | 2 |
| 3.Descriptive Statistics | 2 |
| 4.t-test, Analysis of Variance | 2 |
| 5.$\chi^2$ Test | 2 |
| 6.Nonparametric Test | 2 |
| 7.Linear Regression and Correlation | 2 |
| 8.Statistical Graph | 2 |

### Participants

To study the satisfaction of undergraduates with their curriculum, we administered questionnaire surveys to medical undergraduates at three university medical schools in Guangdong Province (Jinan University, South Medical University, and Guangdong Pharmaceutical University). The participating students were randomly and anonymously chosen from junior medical students at the above-mentioned three universities, and all students completed the medical statistics course during the prior semester. In total, 502 completed questionnaires were eventually received from the 522 issued questionnaires (the response rate of the questionnaire is 96.1%). The respondents included 206 (41.0%) male students and 296 (59.0%) female students; The students were aged between 18-25 (21.6±0.99) years.

To evaluate the teaching effectiveness of a new teaching approach (PDTR), we compared the examination scores of medical graduates who previously studied medical statistics (before 2017) with those of students who were taught using PDTR at Jinan University. In addition, we collected feedback from students who had completed the PDTR course.

## Results

### Investigation of the current teaching status of medical statistics in Chinese higher education

As mentioned in the introduction, medical students at medical schools in China would study almost every chapter of statistical analysis including basic theory, mathematical principles, formula derivation. Concerning the learning effectiveness of the current medical statistics contents, we administered a questionnaire survey to students attending three medical universities/schools in Guangdong Province (Jinan University, South Medical University, and Guangdong Pharmaceutical University). In the survey investigating the students’ favourite aspects of medical statistics, we discovered that 430 and 362 students prefer to study statistical analysis methods (85.7%) and software applications (72.1%); 190 students enjoyed studying about the rationale of medical statistics, while mathematical principles (17.9%) and formula derivation (22.5%) were disliked by 113 and 90 students respectively (Fig. 1). Meanwhile, in the analysis of the general view of the students regarding the study of medical statistics, 231 and 241 students considered the currently used medical statistics textbook (46.0%) and teaching approach inappropriate (48.0%) (Fig. 2), suggesting that some changes in the approaches used to teach medical statistics to medical students are needed.

**Fig. 1.**
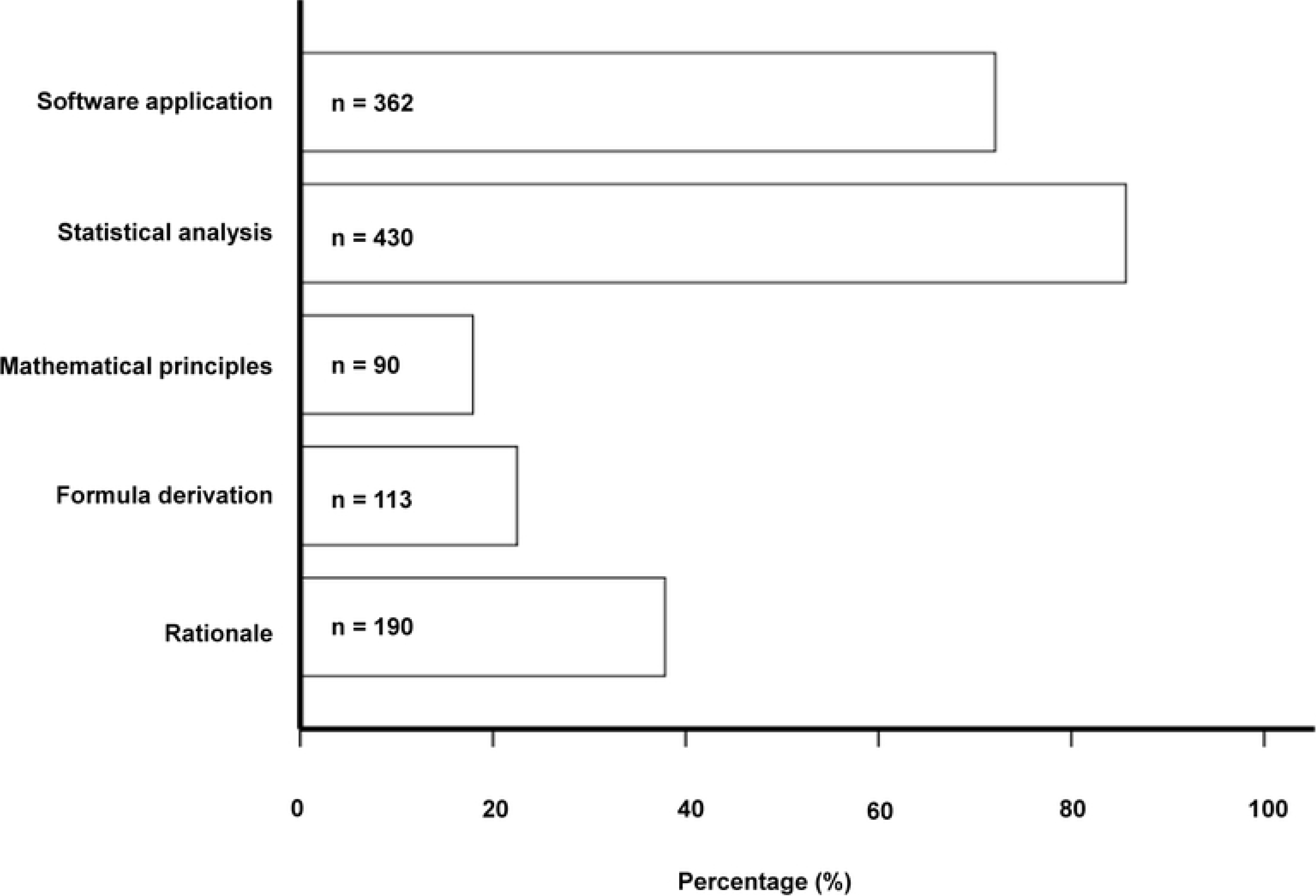
Questionnaire used to survey the students’ favorite content while learning medical statistics among 502 undergraduates attending three medical schools in Guangzhou. The bar chart shows the percentages of 502 medical undergraduate interviewees’ favorite contents among the use of software, rationale, mathematic principles, formula derivation, and statistical analysis while learning medical statistics at Jinan University, Guangdong Pharmaceutical University and Southern Medical University.

**Fig. 2.**
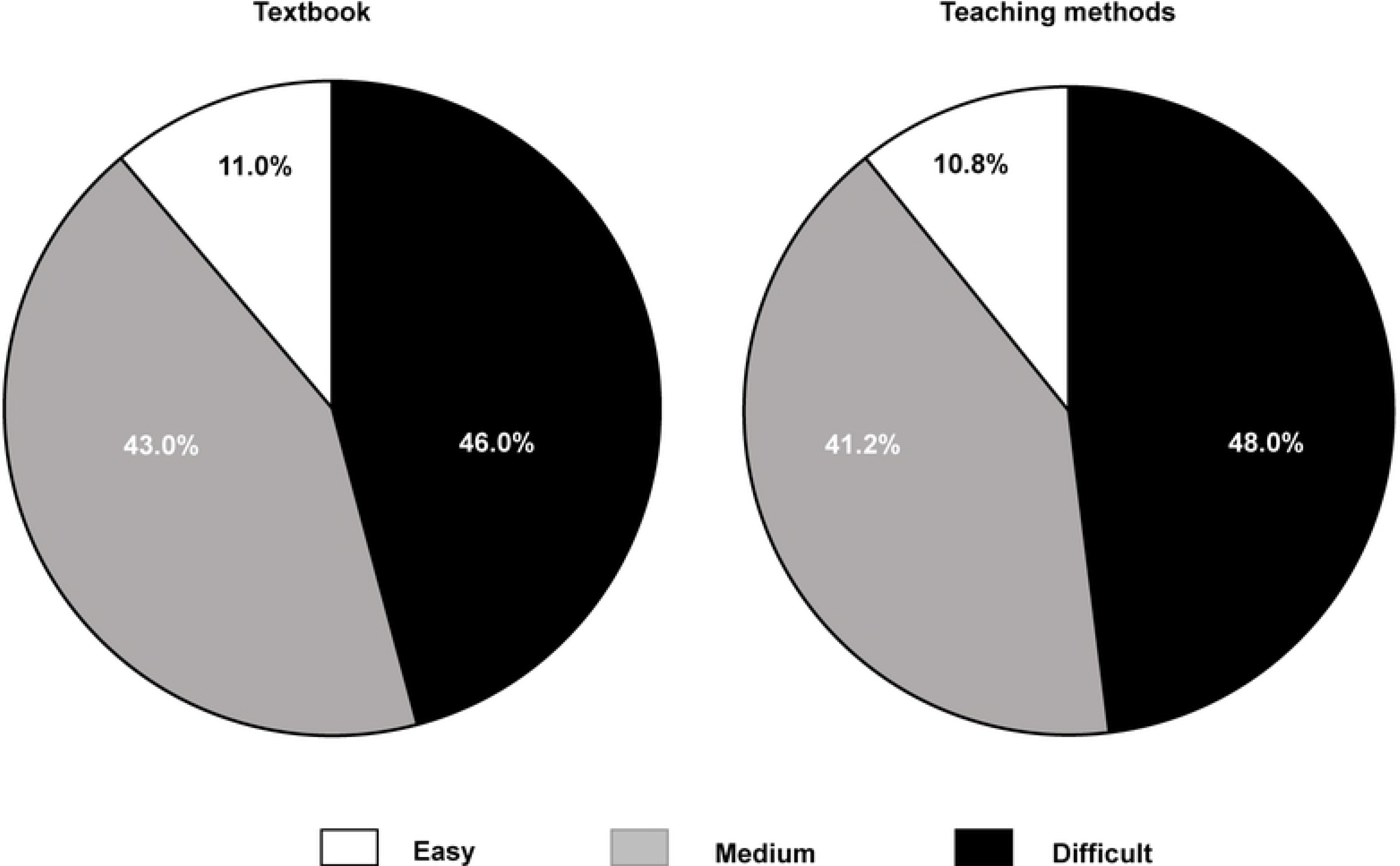
Questionnaire used to survey the students’ reactions to the textbooks and teaching approaches currently used in medical statistics courses among 502 undergraduates attending three medical schools in Guangzhou. The pie chart shows the percentages of 502 medical undergraduate interviewees’ attitudes towards the current textbooks and teaching methods while learning medical statistics at Jinan University, Guangdong Pharmaceutical University and Southern Medical University.

### <u>P</u>urpose, <u>D</u>atabase, <u>T</u>ype of variable, and <u>R</u>elationship between variables (PDTR)

Based on the above-mentioned investigation of the current teaching of medical statistics, most medical students consider it very difficult to understand and grasp current medical statistics. To overcome the difficulties, a new, simpler teaching approach, i.e., PDTR, was developed. The PDTR syllabus is showed in Table 3. Our teaching objective is to train medical students to have statistical thinking, and master basic methods of sorting, summarizing and analyzing medical statistical data. Furthermore, medical students should have basic statistical knowledge and skills for doing scientific research, reading medical literatures and other medical courses. They would learn how to use statistical software, but not limited to SPSS, in other word, they are able to choose any statistical software which is the most appropriate. Totally, this course takes 18 class hours. In addition, this approach focuses on the following four key points:

1. Specifying purpose of analyses: The purpose of an analysis determines the research design, factors, indicators, etc. Statistics is a discipline which is about the “relational data” between influential and outcome variables, students should understand which variable is the influencing variable and which variable is outcome variable before doing statistical analysis. Therefore, the purpose is a fundamental step in statistical analyses. Statistical analyses can be correctly carried out only if the purpose of the analyses is specified.
2. Creating a statistical database: A database refers to a two-dimensional form of orderly recorded observations from different observation indicators based on the research objects. Except for the first row of the two-dimensional table, the remaining rows represent all indicators of an observed object, and each column represents the value of the object of the observed indicator. The database that meets the requirements can make the research indicators of the observed objects clear at a glance, so that the research ideas are specified, and the relevant statistical software can be directly used for analysis and calculation.
3. Identifying the type of variables: The variable types are divided into the influence variable (independent variable) and the outcome variable (dependent variable). The outcome variable is the focus of attention and is the variable we seek to understand. The outcome variable is the result of the change in the influence variable. The influence variables influence the outcome variables. Assessing the size and strength of the influence of one or more influence variables on the outcome variables of interest is a core issue that is emphasized throughout this method. There are four types of variables: numerical variables which are known as quantitative variables, refer to indicators that can be measured by quantitative methods and have numerical values, heights or degrees. Variable values generally have units of measurement might have decimal points (i.e., height and weight); categorical variables, also known as qualitative variables, refer to an attribute classification or feature classification indicator that can be determined by qualitative methods, and are divided into: nominal categorical variables (i.e., place of birth and test group), ordinal categorical variable (i.e., school grade) and binary variables (have only two possible values, such as whether a patient survives or dies).
4. Selecting and performing the appropriate statistical methods: Statistical methods should be selected based on the research purpose and variable types (confirming the relationship between the variables). Since four subtypes of both the outcome variable and influence variable exist, we could obtain a total of sixteen groups of inter-combinations, i.e., at least sixteen types of statistical analysis methods could be chosen, and these methods are presented in detail in Table 4. For instance, if we aim to compare two proportions, i.e., analyse the relationship between two binary variables, a *χ*^2^ test for a 2×2 contingency table should be selected. Alternatively, to analyse the relationship between a numerical variable and an ordinal categorical variable, we should select Spearman rank correlation according to Table 4. If we consider the transforming types of a variable, i.e., numerical variable → ordinal categorical variable → nominal categorical variable → binary variable, many more methods for selection exist.

**Table 3.** Key points and class hour allocation for PDTR

| **Chapter** | **Teaching hour** |
| --- | --- |
| 1. Introduction | 2 |
| 2. How to specify the purpose of analyses | 3 |
| 3. How to create a statistical database | 3 |
| 4. How to identify the variable types | 3 |
| 5. Select and perform the appropriate statistical methods | 4 |
| 6. Application of software | 3 |

**Table 4.**
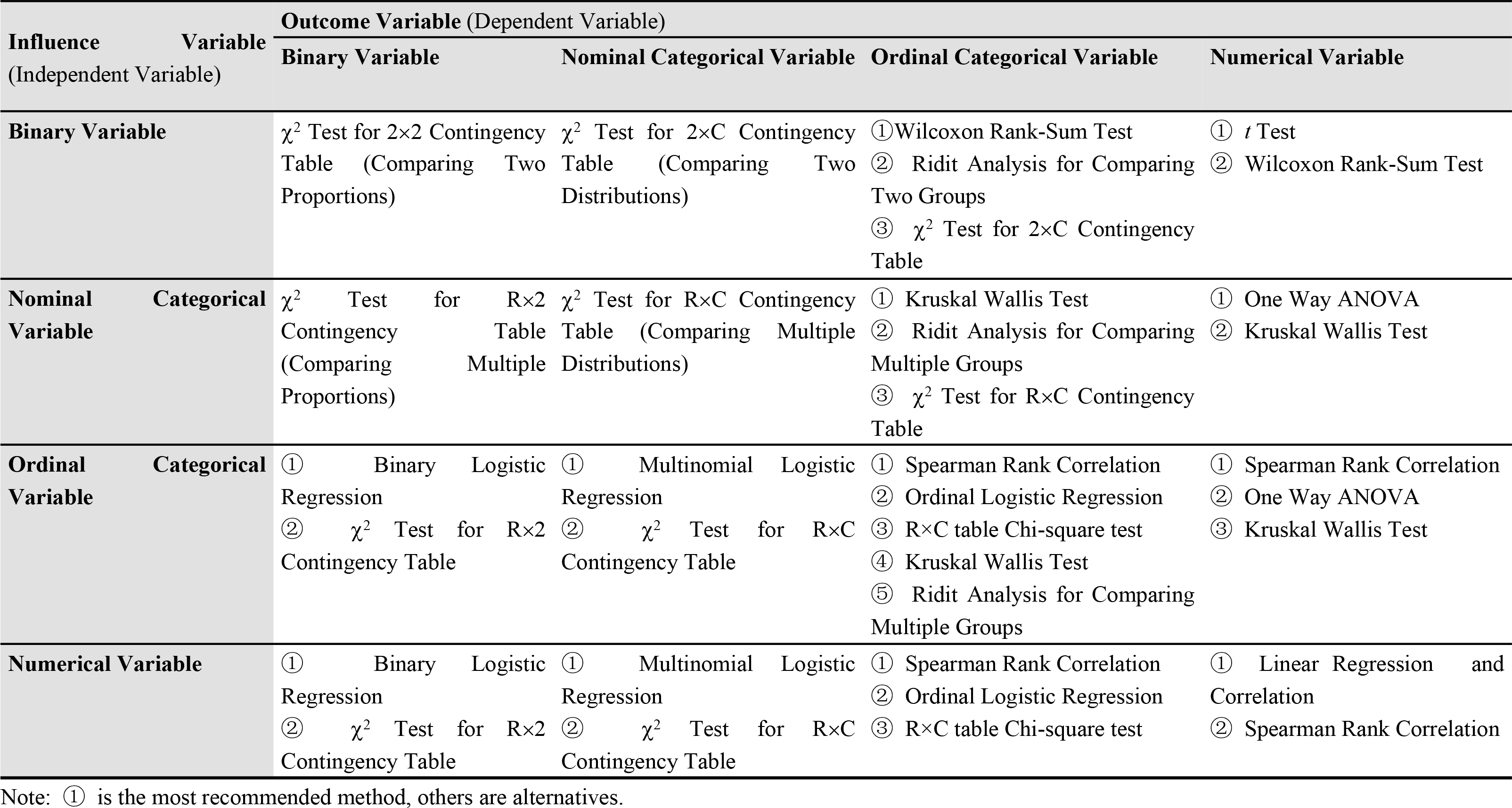
Statistical analysis methods of two variables (PDTR)

### Pilot course “Applied Medical Statistics” at Jinan University Medical School

The new teaching approach PDTR is also named Applied Medical Statistics. Moreover, the pilot course was first offered at Jinan University Medical College to undergraduates and graduates. The participating students were randomly and anonymously chosen from medical students at the Jinan University School of Medicine who have not yet studied medical statistics. The participants included 141 (46.8%) male students and 296 (53.2%) female students; Students were aged between 18-26 (22.1±0.96).

Compared with traditional methods of teaching medical statistics, PDTR is a novel, simple, practical, inflexible and effective approach to teaching medical statistics. Medical students, especially those in clinical medicine, had heavy course load, while the PDTR course could save medical students’ studying time on medical statistics. Students could master basic statistics knowledge quickly and effectively. Moreover, the syllabus can be adjusted based on students’ requirements. Thus, medical students could have more time to practice, such as doing statistical exercise, reading literature and using statistical software for analyzing.

### PDTR application enhanced the learning outcome of the medical students studying medical statistics

To evaluate the effect of PDTR, we carried out examinations with similar difficulty between the graduates of Jinan University Medical School, who had completed the medical statistics course with PDTR or a traditional class (Fig. 3). Meanwhile, qualitative feedback was obtained from the students who had completed the PDTR class. We summarized key points from their feedback (Fig. 4A).

**Fig. 3.**
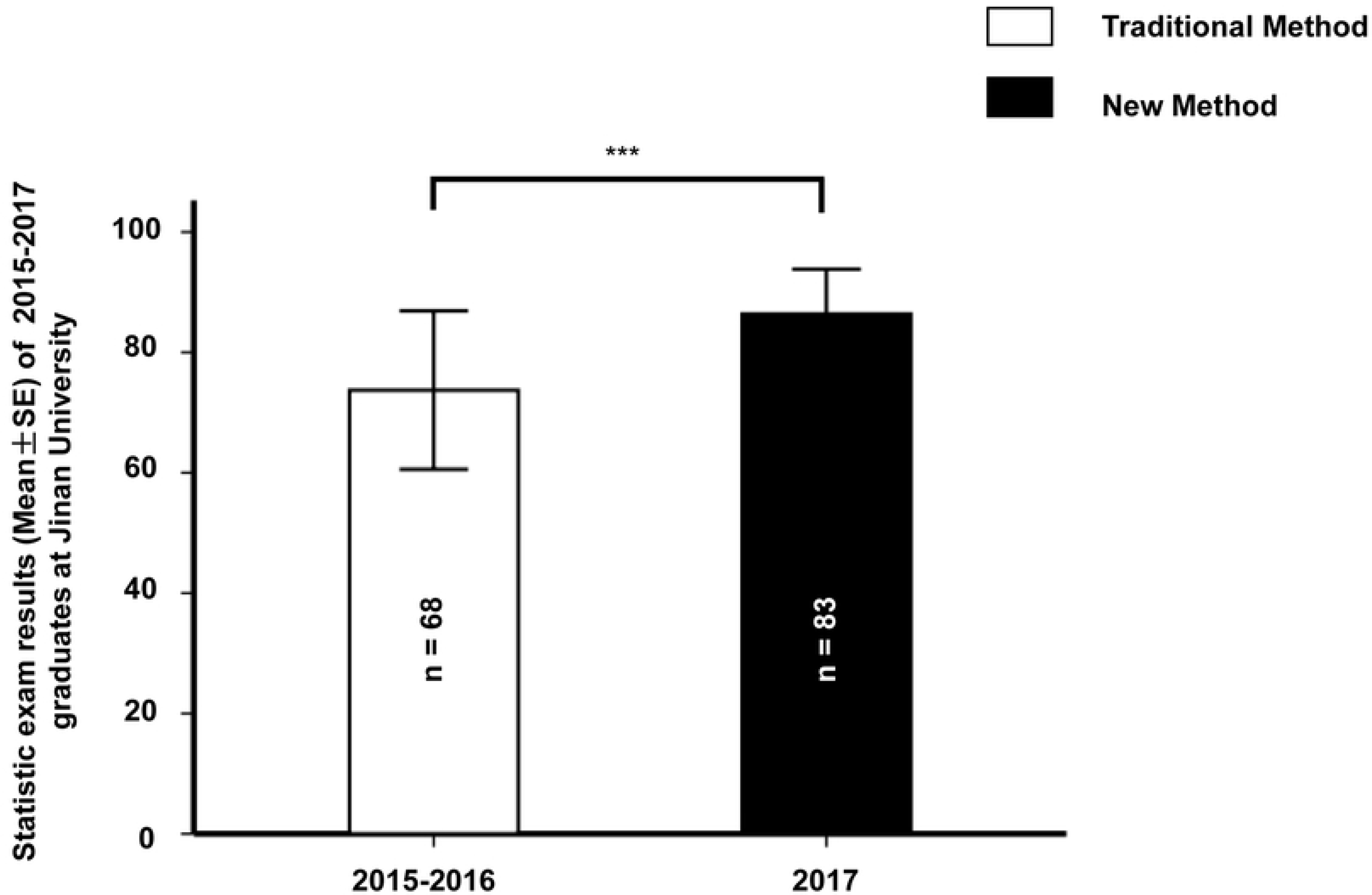
Comparison of the graduates’ examination scores on medical statistics between the traditional and PDTR groups. The bar chart shows a comparison of the examination scores on medical statistics between students who graduated in 2015-2016 (taught using the traditional approach) and students who graduates in 2017 (taught using PDTR at Jinan University Medical School).

**Fig. 4.**
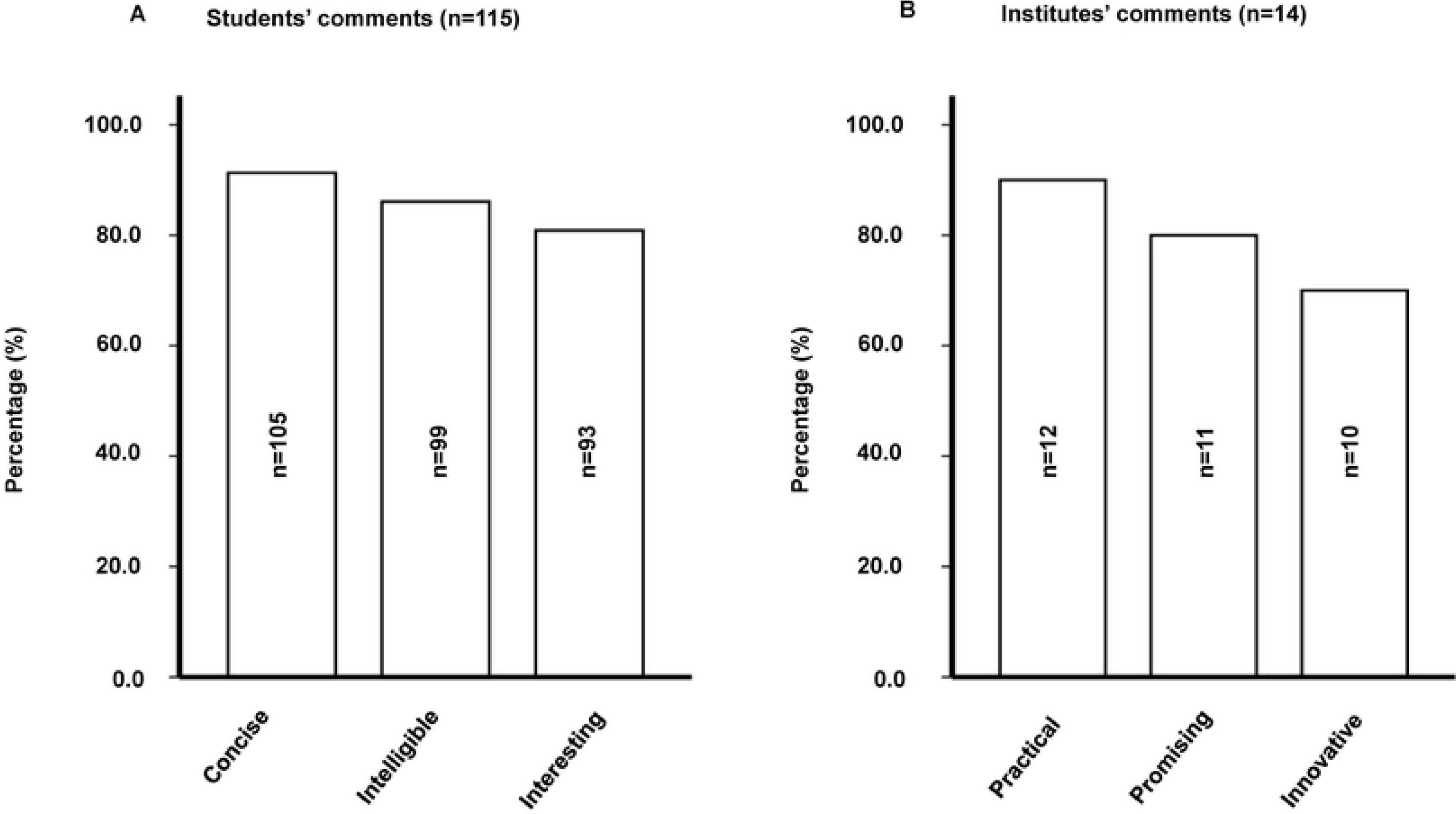
Feedback received from students regarding the performance of PDTR at Jinan University Medical School and other institutes. The bar chart shows the percentages of the participating graduates’ recognitions of PDTR as concise, intelligible and interesting (A); the percentages of institutes’ (at which Dr. Chen taught medical statistics using PDTR) recognitions of PDTR as practical, promising and innovative (B).

The results clearly demonstrated that the examination scores of the medical graduates taught with PDTR were significantly higher than those of the students who were taught with the traditional teaching approach (86.39±3.72 vs 73.72±6.58, *P* < 0.001), suggesting that PDTR indeed improved the learning outcomes of medical graduates completing a medical statistics course.

In addition to Jinan University, the PDTR pilot course was used by an instructor, i.e., Dr. Chen, who teaches medical statistics courses in many other institutes and universities. The key points of feedback were shown in Fig 4. Moreover, the feedback provided by the students regarding the PDTR course in detail was as follows: “I had already studied medical statistics before listening to Dr. Chen’s PDTR course. My impression used to be that medical statistics was boring and elusive. However, PDTR seems to be a completely new subject since it totally changed my traditional idea of medical statistics. I think that it was mainly because PDTR emphasized how to apply practical plans of statistics in daily professional work rather than focusing on teaching mathematical formulas, which greatly stimulates students’ interest in learning statistics. Hence, most of our medical students and physicians do not feel that the course is very difficult with PDTR even with the lack of a strong mathematics background. Anyway, I do think that the pilot course using PDTR is suitable for our medical students and physicians to master and apply medical statistics in professional work.”

## Discussion

As a branch of statistics, medical statistics refers to the application of statistical analysis methods for summarizing, collecting and interpreting data in medical practice and research. Due to the fast development of medicine and health sciences, medical statistics courses are progressively increasingly taught as an integral part of the medical curriculum in all medical schools. Unfortunately, some students did not perceive the importance of medical statistics while studying this topic at school, but they did recognize the value of statistics once they initiated their professional careers (2, 10). However, students frequently dislike studying medical statistics, in turn leading to poor learning outcomes in medical statistics (7). One of the important reasons is that medical students do not have a strong background in mathematics and they have heavy course load. Furthermore, traditional didactic teaching neither activates students’ motivation to learn statistics nor meets their requirements for statistics. Thus, improving how medical statistics are taught to undergraduates and graduates is absolutely necessary to ensure that medical students use statistics appropriately in their professional career.

Actually, how to best teach medical statistics is an ongoing debate (11, 12). For example, Evans et al. proposed that both statisticians and medical colleagues should participate in teams in editing medical statistics curricula to enhance the understanding of the requirements of statistics for medical students/physicians by improving the contents of the teaching material and teaching methods (11). In redesigning the syllabus and the way medical statistics are taught, we sought to make the students’ learning experience more relevant to their eventual medical practice and ensure the students learn the teaching contents(13, 14). In this study, we were also interested in Chinese medical students’ precise perceptions of the currently widely used medical statistics curriculum in China (Table 1). The survey respondents were randomly chosen from three medical schools/universities since these institutions are representative of Chinese medical schools/universities. South Medical University is a typical traditional Chinese medical university, Guangdong Pharmaceutical University represents a medical school at a life science university, and Jinan University is an integrated university including medical schools that recruit medical students globally. Despite the inclusion of different universities/schools, the medical students’ responses regarding the teaching of medical statistics were amazingly consistent. The findings are generally similar to those reported by Hannigan et al. (6), who found that students had negative attitudes and were stressed and discouraged about the current methods used to teach medical statistics, and the students even expressed that they would not choose the course of medical statistics if they had an option. Importantly, we exerted our best efforts to eliminate other possible influencing factors, such as gender. Although a difference in the perception of the difficulty of a medical statistics course between female and male students has been reported (15), we did not find a significant difference in this study; therefore, the gender factor was not particularly mentioned.

Of course, this lack of a difference is partially due to the weakness in the medical students’ mathematics background. However, the often mentioned issue of whether statistics should be regarded as a sub-field of mathematics or an independent discipline emerges (16). From the statistics application perspective, students do not need a strong background in mathematics since mathematics does not directly contribute or necessarily translate to students’ performance in statistics. Statistics, especially medical statistics, should regress to its initial role of analysing variability in data production in the medical profession and research instead of excessively emphasizing the numbers in mathematical statistics significance. The essence of statistics is to make interpretations and critical judgements based on data reported in professional works. In response to these existing issues, Dr. Chen developed a novel approach for teaching medical statistics, i.e., PDTR, which converts the traditional didactic teaching of medical statistics into a simple and compact table application of sixteen types of combinations could directly guide the statistical analysis of two variables, i.e., influence variable and outcome variable (Table 4). Compared to traditional methods of teaching medical statistics, great improvement has been achieved with PDTR, which emphasizes the practical application of statistics and the selection and performance of the appropriate statistical method based on a newly created database that summarizes the relationship between a dependent variable and an independent variable using a table, which certainly reduced the learning difficulty of even students/physicians who do not possess basic knowledge of statistics. It takes less time on learning classroom statistics, and makes students have more choices in using statistical software. Compared to the examination scores on the medical statistics course completed by graduates previously taught using the traditional teaching approach at Jinan University Medical School, the examination scores of the PDTR-taught graduates were significantly higher, implying that the PDTR approach was successful in improving the statistical teaching effectiveness. What’s more, among the graduates at Jinan University and other institutes at which PDTR was employed to teach medical statistics, both the students and the institute organizers fully confirmed the teaching effectiveness of PDTR in terms of its conciseness, practicability, innovativeness, etc. (Fig. 4). As previously mentioned, for the first time, PDTR can effectively improve the learning outcomes of medical statistics, especially of statistical application. Therefore, undoubtedly, widely popularizing this medical statistics teaching approach for medical students is warranted. Of course, further discussion and improvement of PDTR are absolutely required during the practice of this novel teaching approach in the future.

## Conclusion

After precisely surveying the difficult aspects and demands of medical students studying medical statistics at three medical universities/schools in Guangdong Province, we developed a novel model for teaching medical statistics; PDTR was designed to solve the existing problems revealed by the above-mentioned survey of statistics teaching. PDTR emphasizes studying the application of medical statistics using statistical analysis methods involving two variables instead of excessively focusing on statistical theory. The implementation of the PDTR pilot course at Jinan University and other institutes dramatically improved the students’ medical statistics learning outcomes and was praised by the PDTR-taught students and institute organizers.

## Limitations

In this study, we did not limit the particular survey respondents and PDTR pilot course to the same population of students/graduates. In addition, some differences in the students’ mathematical backgrounds might exist among the three surveyed medical universities/schools, although all participants were medical students. The students were combined, rather than divided by each university/school, in the analysis of the trends of the students’ choices on their questionnaires. Furthermore, we did not divide the exchange students into groups based on their interests in learning. Although PDTR is a novel and simple teaching method, it does not cover some advanced statistical methods, such as survival analysis and cluster analysis. Undoubtedly, this approach should be improved based on feedback from students and instructors in the future.

## Ethics approval

The ethics committee of Jinan University carefully considered and approved the project proposal.

## Consent for publication

All authors have seen the manuscript and approved to submit.

## Acknowledgements

The authors would to thank all participants who filled out the questionnaire, took part in the course and provide feedback.

## References

1. Greig JD. The westernization of Chinese medicine and medical education. Hangzhou leads by example. Scottish Medical Journal. 1994;39(4):123–5.

2. Miles S, Price GM, Swift L, Shepstone L, Leinster SJ. Statistics teaching in medical school: opinions of practising doctors. BMC medical education. 2010 Nov 4;10:75. PubMed PMID: 21050444. Pubmed Central PMCID: 2987935.

3. Herman A, Notzer N, Libman Z, Braunstein R, Steinberg DM. Statistical education for medical students--concepts are what remain when the details are forgotten. Statistics in medicine. 2007 Oct 15;26(23):4344–51. PubMed PMID: 17487940.

4. Morris RW. Does EBM offer the best opportunity yet for teaching medical statistics? Statistics in medicine. 2010;21(7):969–77.

5. Zhang Y, Shang L, Wang R, Zhao Q, Li C, Xu Y, et al. Attitudes toward statistics in medical postgraduates: measuring, evaluating and monitoring. BMC medical education. 2012 Nov 23;12:117. PubMed PMID: 23173770. Pubmed Central PMCID: 3533942.

6. Hannigan A, Hegarty AC, McGrath D. Attitudes towards statistics of graduate entry medical students: the role of prior learning experiences. BMC medical education. 2014 Apr 4;14:70. PubMed PMID: 24708762. Pubmed Central PMCID: 4234395.

7. Freeman JV, Collier S, Staniforth D, Smith KJ. Innovations in curriculum design: a multi-disciplinary approach to teaching statistics to undergraduate medical students. BMC medical education. 2008 May 1;8:28. PubMed PMID: 18452599. Pubmed Central PMCID: 2397402.

8. Storrar N, Hope D, Cameron H. Student perspective on outcomes and process - Recommendations for implementing competency-based medical education. Medical teacher. 2018 Mar 20:1–6. PubMed PMID: 29557693.

9. Sheng Y. [Sixty years of study on history of chinese medical education]. Zhonghua yi shi za zhi. 1996;26(3):170–8. PubMed PMID: 11614110.

10. Windish DM, Huot SJ, Green ML. Medicine residents’ understanding of the biostatistics and results in the medical literature. Jama. 2007 Sep 5;298(9):1010–22. PubMed PMID: 17785646.

11. Evans SJ. Statistics for medical students in the 1990’s: how should we approach the future? Statistics in medicine. 1990 Sep;9(9):1069–75; discussion 77-8. PubMed PMID: 2244079.

12. Clayden AD. Who should teach medical statistics, when, how and where should it be taught? Statistics in medicine. 1990 Sep;9(9):1031–7; discussion 9-44. PubMed PMID: 2244076.

13. Sterne JAC. Teaching hypothesis tests – time for significant change? Statistics in medicine. 2002;21(7):985–94.

14. Astin J, Jenkins T, Moore L. Medical students’ perspective on the teaching of medical statistics in the undergraduate medical curriculum. Statistics in medicine. 2002 Apr 15;21(7):1003–6; discussion 7. PubMed PMID: 11921009.

15. Hilton SC, Schau C, Olsen JA. Survey of attitudes toward statistics: Factor structure invariance by gender and by administration time. Struct Equ Modeling. 2004;11(1):92–109. PubMed PMID: WOS:000221781000007. English.

16. The challenge of developing statistical literacy, reasoning and thinking. J Res Math Educ. 2006 Mar;37(2):151–. PubMed PMID: WOS:000235758300006. English.

